# EEG frequency tagging evidence of intact social interaction recognition in adults with autism

**DOI:** 10.1101/2022.07.14.500030

**Authors:** Danna Oomen, Emiel Cracco, Marcel Brass, Jan R. Wiersema

**Author notes:** **Author Note:** Correspondence concerning this article should be addressed to Danna Oomen, Department of Experimental Clinical and Health Psychology, Ghent University, Henri Dunantlaan 2, B-9000, Ghent, Belgium.

## Abstract

To explain the social difficulties in autism, a large amount of research has been conducted on the neural correlates of social perception. However, this research has mostly used basic social stimuli (e.g. eyes, faces, hands, single agent), not resembling the complexity of what we encounter in our daily social lives, and as such, the situations people with autism experience difficulties in. A more complex stimulus that we do come across often and is also highly relevant for social functioning is that of third-party social interactions. Here, we investigated if individuals with and without autism process third-party social interactions differently. More specifically, we measured neural responses to social scenes depicting either social interaction or not with an electroencephalogram (EEG) frequency tagging task and compared these responses between adults with and without autism (*N* = 61). The results revealed an enhanced response to social scenes with interaction, replicating previous findings in a neurotypical sample (Oomen et al., 2022). Crucially, this effect was found in both groups with no difference between them. This suggest that social interaction recognition is not anomalous in adults with autism and cannot explain the social difficulties adults with autism experience.

**Lay abstract:** People with autism have social difficulties and are thought to experience the world differently. To better understand these differences, research has studied how the brain of people with and without autism processes social stimuli. However, this research has mostly used basic social stimuli (e.g. eyes, faces, hands, and single agents). Such stimuli do not resemble the complexity of daily life, where we typically do not come across isolated body parts, but instead have to make sense of complex social scenes with multiple people. To do so, it is imperative that we are able to recognize social interaction. Hence, if social interaction processing is anomalous, this could have pervasive consequences for social functioning more generally. Here, we used brain imaging to test if adults with autism process social interaction scenes differently than adults without autism. In line with previous findings from a neurotypical sample (Oomen et al. (2022), we found that social scenes depicting interaction elicited stronger brain responses than social scenes not depicting interaction. Crucially, this effect was found in both groups with no difference between them. These findings suggest that the fundamental process of social interaction recognition is not anomalous in adults with autism.

## 1. Introduction

The social world can be challenging for people with autism spectrum disorder^1^ (ASD; henceforth ‘autism’), who experience difficulties in social interaction and communication, including non-verbal communicative behaviour (American Psychiatric Association, 2013). In order to explain the social difficulties associated with this neurodevelopmental disorder, much research has been conducted on the neural correlates of social stimuli processing. This research typically uses simple stimuli such as a single agent or even isolated body parts like faces, eyes or hands (single agent: e.g. Nijhof et al., 2018; faces: e.g. Kang et al., 2018; eyes: Holt et al., 2014 hands: e.g. Raymaekers et al., 2009, Okamoto et al., 2018), often stripped from any contextual background information (i.e. put against a blank background). Although this has the advantage of increasing experimental control, it does not resemble the complexity of what we encounter in real life, a problem that is especially relevant in autism research, where it has become increasingly clear that anomalies mainly exist for processing more complex social stimuli (Dziobek et al., 2006; Golan et al., 2008; Heavey et al., 2000; Roeyers et al., 2001).

Considering this research, a particularly relevant type of stimuli to understand the social difficulties in autism are third-party social interactions (Quadflieg & Koldewyn, 2017). Indeed, not only does processing social interaction require complex cognitive functioning (Isik et al., 2020), it is also highly relevant for social functioning. For example, research has shown that our impressions of people are strongly shaped by how they interact with others (Costanzo & Archer, 1989; Cowell & Decety, 2015; Mast & Hall, 2004; Sinke et al., 2010), which in turn determines our attitudes towards them (Quadflieg & Penton-Voak, 2017) and guides our own actions (Christ et al., 2014). Social interaction recognition also, quite literally, navigates us through the social world, helping us to not break up interactions when walking through a crowded environment (Efran & Cheyne, 1973; Knowles, 2015). Hence, if the fundamental process of social interaction recognition is anomalous in autism, this could have cascading effects on how people with autism process, experience, and take part in the social world, potentially explaining part of why this differs from people without autism.

Existing studies on social interaction processing in autism have used behavioural methods (Liu et al., 2018; van Boxtel et al., 2017; von der Lühe et al., 2016). First, van Boxtel et al. (2017) found that neurotypical adults who score relatively high on autism symptomatology showed a reduced ability to differentiate between interactive and non-interactive actions. Similarly, Liu et al. (2018) showed that motion sequences that depict agents in social interaction tend to be perceived as shorter in duration by neurotypicals, but that this effect was negatively correlated with autism symptomatology. Finally, von der Lühe et al. (2016) found that adults with autism show diminished interpersonal predictive coding. That is, their autism sample made less use of the actions of one agent to predict the actions of another agent. Taken together, these behavioural studies suggest that individuals with autism process social interactions differently than individuals without autism.

Importantly, however, this could have two reasons: 1) it could mean that individuals with autism have difficulties to recognize social interaction, but 2) could also mean that they are perfectly able to recognize social interactions, but do not extract the same information from those interactions and/or use this information similarly. To clarify the latter, behavioural studies necessarily impose a task. As a result, when performance on that task differs, this does not necessarily mean that participants with autism did not recognize social interaction, but could also mean that they used the information extracted from those interactions differently to perform the task. For example, in the study by van van Boxtel et al. (2017), participants had to indicate the degree of perceived interaction of interacting and non-interacting pairs. However, differences in such a task do not necessarily reflect a perceptual bias but could also reflect a response bias. For example, individuals with autism may have a higher threshold to explicitly label something as interaction. To test if individuals with autism truly have difficulties to recognize social interaction, we therefore have to measure spontaneous social interaction processing without an explicit task.

Oomen et al. (2022) recently developed and validated an electroencephalogram (EEG) frequency tagging task that can be used to do this. More specifically, in separate blocks, they presented stimuli of social scenes depicting either two interacting agents or non-interacting agents at a fixed frequency. This produced a neural response at exactly that frequency, which revealed an enhanced response over predominantly right lateralized occipitoparietal electrodes, to social scenes with social interaction compared to social scenes without social interaction. By presenting the stimuli at a fixed rate the brain response is restricted to a narrow frequency band, which gives the technique the advantage of being largely resistant to noise and therefore providing a high signal to noise ratio (Norcia et al., 2015a; Retter & Rossion, 2016). Most crucially however, the method applied by Oomen et al. (2022) provides an objective measure of spontaneous social interaction recognition as no explicit task is required. Hence, in the current study, we applied the same task and technique in a sample of adults with and without autism (neurotypical adults) to directly investigate whether third-party social interaction recognition differs between these groups. We expected to replicate the finding of Oomen et al. (2022) and to observe a stronger brain response for interaction than for non-interaction scenes in neurotypical adults. Furthermore, we hypothesized that if the fundamental process of social interaction recognition would be anomalous in autism, that this effect would be absent or diminished in individuals with autism.

### Open science statement

Our hypotheses, study design, and data-analyses plan were preregistered (https://aspredicted.org/blind.php?x=J6G_DR9). Data and analyses can be found on the Open Science Framework (https://osf.io/qt7z3/?view_only=0da455e567d742ce9dc0b4209238372d)

### Community involvement statement

We want to be respectful in the way we refer to our participants with a diagnosis of autism spectrum disorder. As most published academic research on this topic is from the UK or USA, and terminology preferences may differ internationally (taking in account cultural and linguistic contexts), we let our participants inform us of their preferences. Sixteen participants preferred ‘with autism’, 11 participants had no opinion, 2 participants preferred ‘with ASD’, 2 participants preferred ‘on the spectrum’, and 1 participant preferred ‘autistic’. All participants gave consent for us to use the terminology most agreed on. Here, we therefore use ‘person with autism’ to respect the preference of the autism sample described in this study.

## 2. Methods

### 2.1 Participants

We tested 32 adults with autism (autism group) and 32 neurotypical adults (control group). All 64 participants reported to have normal or corrected-to-normal vision, no neurological condition, and sufficient knowledge of the Dutch language. Participants in the control group reported no known psychiatric condition and had a T-score of 60 or lower on the Social Responsiveness Scale – adult version (SRS-A; Constantino, 2002; Dutch version: Noens et al., 2012). Participants in the autism group had a formal diagnosis of autism^2^, and were only included if they had a T-score of 61 or higher on the SRS-A (Constantino, 2002; Dutch version: Noens et al., 2012). The SRS-A is a 64-item questionnaire used to measure autism traits with a recommended cut-off of 61. Lastly, participants for both groups were only included if they had an IQ score of 85 or above (i.e. average or above). IQ scores were retrieved from the lab database in case the participant had previously participated in research of the lab. Alternatively, reports were obtained if an IQ test had been administered elsewhere (e.g. during the diagnostic procedure). If a participant’s IQ score was not known yet, or acquired before adulthood, an estimate was obtained after the test session. For this, we administered four subtests of the WAIS-IV-NL (i.e. Block design, Vocabulary, Matrix reasoning, Similarities; Wechsler, 2008), corresponding to the WASI-II short four-subtest form (Wechsler, 2008), in order to obtain WASI-II scores but with Flemish norms. Three participants were excluded overall. One participant was excluded from the autism group due to an SRS score below the cut-off. Two participants were excluded from the control group: one due to an IQ score lower than 85, the other due to bad data quality. In line with our aspired, preregistered sample size, the final sample thus consisted of 31 adults with autism and 30 neurotypical adults. Note that although we did not preregister to exclude participants based on SRS-A and IQ scores, this is in line with previous research in autism (e.g. Goris et al., 2018; Vettori et al., 2019). Importantly, including the two participants whose exclusion was not based on our preregistered exclusion criteria did not change any of the results reported here.

Groups were matched on age, gender, and IQ (see Table 1) and as expected differed on the SRS-A as well as on the autism-spectrum quotient (AQ) and Toronto Alexithymia Scale (TAS-20). The AQ is another commonly used questionnaire to measure autism traits through 50 self-report items (Baron-Cohen et al., 2001; Dutch version: Hoekstra et al., 2008). The TAS-20 is a 20-item self-report to measure alexithymia (Bagby et al., 1994; Dutch version: Trijsburg et al., 1997), which is relatively common in autism (Kinnaird et al., 2019). The AQ and TAS-20 were administered to further describe our sample and, together with the SRS, to conduct exploratory correlational analyses. Note that the AQ and SRS-A scores were comparable to other studies that included adults with and without autism (e.g. Goris et al., 2022; Nijhof et al., 2018). The experimental protocol was approved by the local ethics committee of the Faculty of Psychology and Educational Sciences of Ghent University (EC/2020/122), and written informed consent was obtained from all participants before the start of the study. Participants were compensated for their time.

**Table 1.** Participant characteristics

|  | autism<br>*M (SD)* | control<br>*M (SD)* | *F (p)* |
| --- | --- | --- | --- |
| Age | 35.00 (7.30) | 34.23 (7.70) |  |
| Gender |  |  |  |
| Female | 11 | 11 |  |
| Male | 18 | 19 |  |
| Non-binary | 2 | 0 |  |
| Ethnicity |  |  |  |
| White | 30 | 28 |  |
| Asian | 0 | 1 |  |
| Mixed | 1 | 1 |  |
| IQ | 110.45 (11.12) | 106.80 (7.92) |  |
| SRS-A T-score | 75.16 (8.32) | 48.37 (6.90) | 186.80 (< .001) |
| AQ | 35.35 (5.72) | 14.40 (7.01) | 164.14 (< .001) |
| TAS-20 | 58.09 (9.92) | 42.00 (8.89) | 49.02 (< .001) |

*Note.* Autism group $n = 31$ ; control group $n = 30$ ; SRS-A = social responsiveness scale – adults; AQ = autism-spectrum quotient; TAS-20: Toronto Alexithymia Scale – 20 items

### 2.2 Frequency Tagging Task and Procedure

Participants were seated in a Faraday cage circa 80-100 cm from a 24-inch computer screen with a refresh rate of 60 Hz. Participants filled in the SRS-A and AQ questionnaires, after which they completed two frequency tagging tasks intermitted by three minutes of resting state EEG. The order of the two tasks was counterbalanced across participants. Only the task that answers our current research question is described here.

The task was identical to the one described by Oomen et al. (2022). The task included four types of images: 36 images that depicted social interaction, 36 images that depicted no social interaction, and to control for potential low-level differences between the two image types, the scrambled versions of both image types. The scrambled images were created by scrambling the images into a 10 × 10 grid. All images included exactly two agents. Other than that, the images within the interaction and non-interaction categories differed greatly from each other in terms agent configuration (e.g. facing or not facing), activity (e.g. talking or playing), and/or contextual background (e.g. supermarket, school), to resemble the complexity of real-life situations. To increase experimental control, the images were black-and-white line drawings, and agents and objects were free from distractors (e.g. shadows, patterns on clothing). Figure 1 shows one example image of each stimulus type. The two types of images were balanced for perceived emotional valance and perceived intensity of the feelings experienced by the agents.

**Figure 1.**
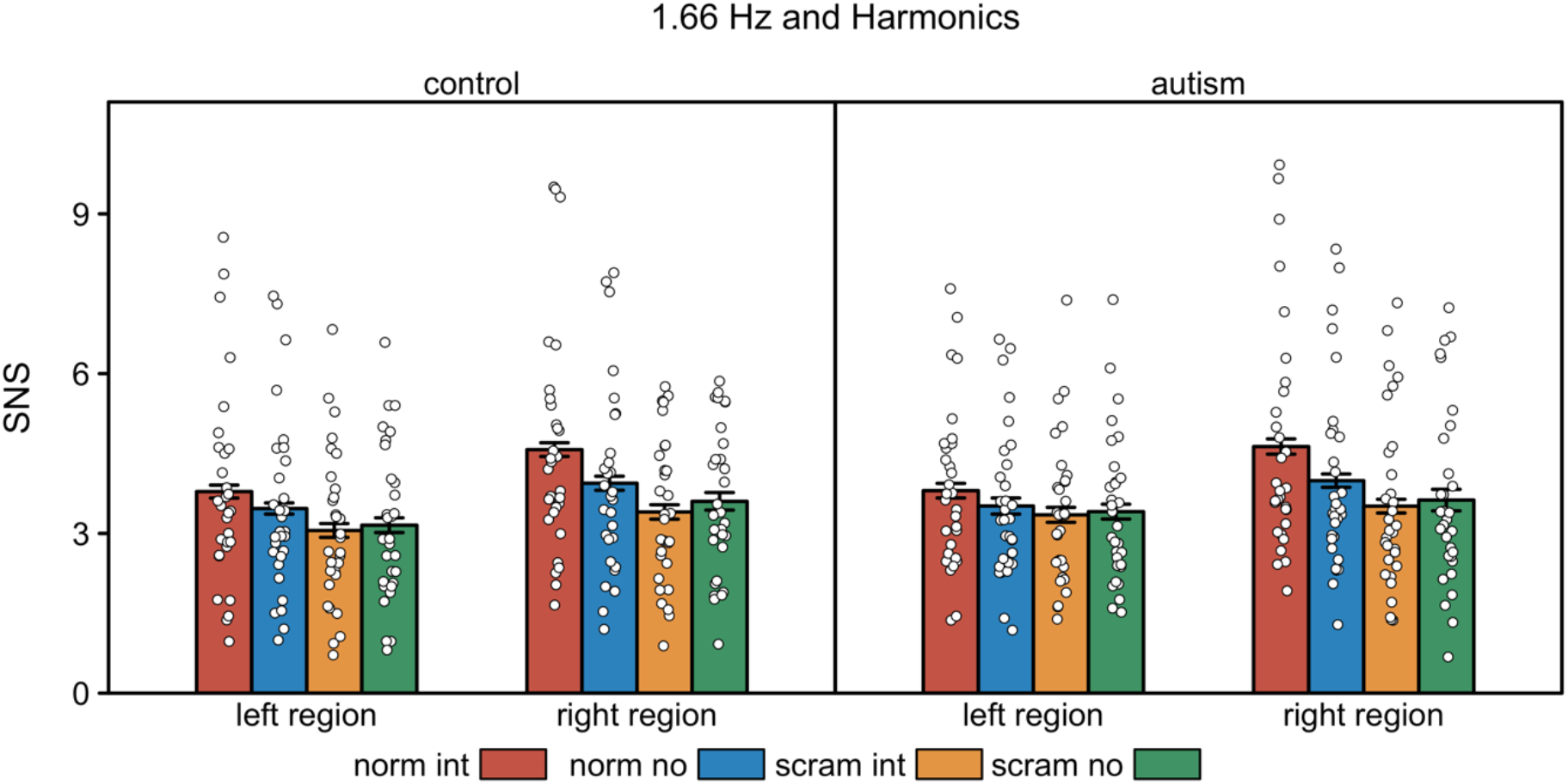
Signal to noise-subtracted amplitudes (SNS) per condition and region and for the two groups separately. Error bars represent errors of the mean (SEMs) corrected for within-subject designs (Morey, 2008).

Images were presented at the centre of the screen using sinusoidal contrast modulation at a presentation rate of 1.66 Hz (600 ms). The four types of images were presented block-wise in blocks of 110 images, with four blocks per category, presented in random order. Images were drawn randomly from their respective categories, never repeating the same image back-to-back. A block started and ended with a 3 s fade in and out (image transparency from 0 to 100% and 100% to 0%). The duration of the task was approximately 18 minutes. The task was programmed in PsychoPy3 (Peirce et al., 2019).

Before the start of the task, participants were told that they would see images of social interaction, images without social interaction, and scrambled images. This information was given together with two example images of the interaction and no-interaction categories that were not included in the actual task. In line with Oomen et al. (2022), we were open about the included stimulus categories to reduce variability regarding the timepoint at which participants recognized them. Participants had two tasks, of which the sole aim was to encourage a constant level of attention. One task was to press the spacebar as fast and accurately as possible whenever the black fixation cross changed to red (400ms), 3 to 6 times per block. Detection was high overall (96% on average). A mixed ANOVA with Interaction Type (no interaction, interaction) and Stimulus Type (scrambled, normal) as within-subject factors and Group (control, autism) as between-subject factor revealed no main or interaction effects on detection rate (all *p*s ≥ .078). The second task was a memory task. Participants were instructed to pay attention to the presented images during the task, so that they would be able to indicate which 4 images (out of 8) had appeared during the task. On average, participants recognized 3.49 out of 4 images. A mixed ANOVA with Interaction type (no interaction, interaction) as within-subject factor and Group (control, autism) as between-subject factor revealed no main effects or interaction effect on recognition (all *p*s ≥ .390).

### 2.3 EEG Recording, Pre-processing, and Analyses

EEG was continuously recorded with 64 Ag/AgCI (active) electrodes that were mounted in an elastic cap (ActiCAP, Munich, Germany), using an ActiCHamp amplifier and BrainVision Recorder software (version 1.21.0402, Brain Products, Gilching, Germany). The sample rate was 100 Hz. Electrodes were positioned mostly according to the 10%-system, with the exception of two electrodes (TP9 and TP10) that were placed at OI1h and OI2h according to the 5%-system to cover a wider area of posterior-occipital activation. Additional bipolar AG/AgCI sintered ring electrodes were placed above and below the left eye to record vertical electro-oculogram (EOG). FT9 and FT10 electrodes were used to record Horizontal EOG. Fz was used as online reference.

Off-line pre-processing of the raw data was done using Letswave 6 (www.letswave.org). First, a fourth-order Butterworth band-pass filter was applied to the data with a low and high cut-off of 0.1 Hz and 100 Hz. The data was then segmented according to the four block types. To remove eye blinks, we computed an ICA matrix for each participant on the merged segmented data sets using the Runica algorithm and a square matrix. ICs of each participant were inspected and the IC-related to eye blinks were manually removed. Next, noisy electrodes were interpolated using three (or two in case of OI1/2h electrodes) neighbouring electrodes. After interpolation, the data was re-referenced to an average reference and Fz was included as a regular electrode. The fade in and out were then cropped from the signal, resulting in 60 s epochs that started at 3 s and ended at 63 s after the onset of a block. Next, epochs within each block type were averaged and a Fast Fourier Transform (FFT) was applied to transform the data of each electrode to normalized (divided by N/2) amplitudes (µV) in the frequency domain. Finally, for the statistical analyses, we computed and exported the signal to noise-subtracted amplitudes (SNS) at each frequency bin by subtracting the average voltage amplitude of the 20 neighbouring bins (10 on each side, excluding the immediately adjacent bin), from the amplitude of the frequencies of interests. For visualization we computed the signal-to-noise ratio (SNR). The SNR computation was identical to the SNS computation, except that division was used instead of subtraction.

Statistical analyses on the SNS data were conducted in R (R Core Team, 2017). Frequency tagged brain responses are often not only evident at the frequency of stimulation (*F*) but also across its higher harmonics (2*F*, 3*F*, etc.). Therefore, to accurately capture the evoked brain response, the relevant harmonics should be summed (Retter et al., 2021; Retter & Rossion, 2016). Based on the previous study (Oomen et al., 2022), which was in turn based on visual inspection of pilot data, we included the first 8 harmonics (1.66 Hz, 3.33 Hz, 5.00 Hz, 6.66 Hz, 8.33 Hz, 10.00 Hz, 11.66 Hz, 13.33 Hz). To ensure an unbiased selection of electrodes, independent of condition effects or hypotheses, region of interests were chosen based on visual inspection of the scalp topography across groups and conditions (collapsed localizer approach(Luck & Gaspelin, 2017; see Supplementary Material for the collapsed scalp topography). As the scalp topography matched that of the previous study (Oomen et al., 2022), namely lateral posterior activity, we used the same clusters and electrodes: a right posterior cluster including PO8, PO4, and O2, and a left posterior cluster including the corresponding electrodes on the left hemisphere, namely PO7, PO3, and O1.

On the SNS data, we conducted a mixed ANOVA with Interaction Type (no interaction, interaction), Stimulus Type (scrambled, normal), and Laterality (left, right) as within-subject factors, and Group (control, autism) as between-subject factor. Interactions between factors were followed up with two-tailed *t*-test. As pre-registered, we also explored the relationship between the Interaction Type x Stimulus Type interaction effect ([normal interaction - scrambled interaction] - [normal no interaction - scrambled no interaction]) and participants’ scores on the SRS-A, AQ, and TAS-20. For this we performed a Spearman rho correlation. For the correlation, *t*-tests, and *F*-tests, we accompanied the *p*-values with Bayes Factors (BFs). BFs were calculated with a noninformative Jeffreys prior, a Cauchy prior on the standardized effect size, and a default prior (Rouder et al., 2012).

## 3. Results

The repeated measures ANOVA revealed a significant main effect of Interaction Type, Stimulus Type and Laterality. The main effect of Interaction type, *F*(1, 59) = 14.18, *p* < .001, η_p_^2^ = .19, BF_10_ = 6.36E+, revealed a stronger response for the interaction stimuli (*M* = 3.76, *SD* = 1.50) than for the no-interaction stimuli (*M* = 3.59, *SD* = 1.40). The main effect of Stimulus Type, *F*(1, 59) = 16.32, *p* < .001, η_p_^2^ = .22, BF_10_ = 6.67E+3, revealed a stronger response for the normal stimuli (*M* = 3.96, *SD* = 1.65) than for the scrambled stimuli (*M* = 3.39, *SD* =1.35). The main effect of Laterality, *F*(1, 59) = 7.63, *p* = .008, η_p_^2^ = .11, BF_10 =_ 39.77, revealed a stronger response in the right cluster (*M* = 3.91, *SD* = 1.63) than in the left cluster (*M* = 3.45, *SD* = 1.43). There was no main effect of Group, *F*(1, 59) = 0.08, *p* = .777, η_p_^2^ = .00, BF_10 =_ 0.27.

The repeated measures ANOVA further revealed an Interaction Type x Stimulus Type interaction, *F*(1, 59) = 48.18, *p* < .001, η_p_^2^ = .45, BF_10_ = 3.23E+10, and an Interaction Type x Laterality interaction, *F*(1, 59) = 4.83, *p* = .032, η_p_^2^ = .08, BF_10_ = 69.13, which were further qualified by an Interaction Type x Stimulus Type x Laterality three-way interaction, *F*(1, 59) = 12.73, *p* = .001, η_p_^2^ = .18, BF_10_ = 136.99. To follow up on this three-way interaction, we looked at the Interaction Type x Stimulus Type effect separately for the left and right cluster (see Table 2 for means and standard deviations). This revealed that there was an Interaction Type x Stimulus Type interaction for both clusters, but this effect was stronger in the right cluster, *F*(1, 60) = 93.11, *p* < .001, η_p_^2^ = .61, BF_10_ = 9.35E+10, than in the left cluster, *F*(1, 60) = 33.37, *p* < .001, η_p_^2^ = .36, BF_10_ =5.04E+4. In both clusters, the Interaction Type x Stimulus Type effect indicated that responses were stronger for interacting than for non-interacting stimuli when they were presented normal (Left: *t*(60) = 6.13, *p* < .001, *d*_z_ = 0.78, BF_10_ = 1.81E+5; Right: *t*(60) = 9.31, *p* < .001, *d*_z_ = 1.19, BF_10_ = 2.69E+10), but not when they were presented scrambled. Instead, for the scrambled images, a weaker effect in the opposite direction emerged, with stronger responses for non-interacting than for interacting stimuli (Left: *t*(60) = -2.03, *p* = .046, *d*_z_ = 0.26, BF_10_ = 0.95; Right: *t*(60) = -3.81, *p* < .001, *d*_z_ = 0.49, BF_10_ = 72.89).

**Table 2.** Means and standard deviations per stimuli condition

| Stimuli | $M(SD)$ |
| --- | --- |
| Left region |  |
| Normal interaction | 3.80 (1.69) |
| Normal no interaction | 3.49 (1.51) |
| Scrambled interaction | 3.20 (1.39) |
| Scrambled no interaction | 3.28 (1.41) |
| Right region |  |
| Normal interaction | 4.60 (2.08) |
| Normal no interaction | 3.97 (1.73) |
| Scrambled interaction | 3.46 (1.49) |
| Scrambled no interaction | 3.61 (1.53) |

In contrast with our hypothesis, we did *not* find a Group x Interaction Type x Stimulus Type effect, *F*(1, 59) = 0.33, *p* = .565, η_p_^2^ = .01. Supporting this non-significant Group x Interaction type x Stimulus Type effect, a Bayesian test of the 3-way interaction showed moderate evidence for H0 (BF_10_ = 0.30). Similarly, separate tests of the Interaction Type x Stimulus Type effect in the two groups revealed a significant interaction for both the control group, *F*(1, 29) = 45.68, *p* < .001, η_p_^2^ = .61, BF_10_ = 7.74E+4, and the autism group, *F*(1, 30) = 41.00, *p* < .001, η_p_^2^ = .58, BF_10_ = 3.65E+4. None of the other effects not reported here reached significance, all *p* ≥ 125. See Figure 1 for a visualization of the SNS data of all stimuli conditions separately for region and group, Figure 2 for a SNR plot over electrodes of interest per condition per group, and Figure 3 for the topographies per condition per group.

**Figure 2.**
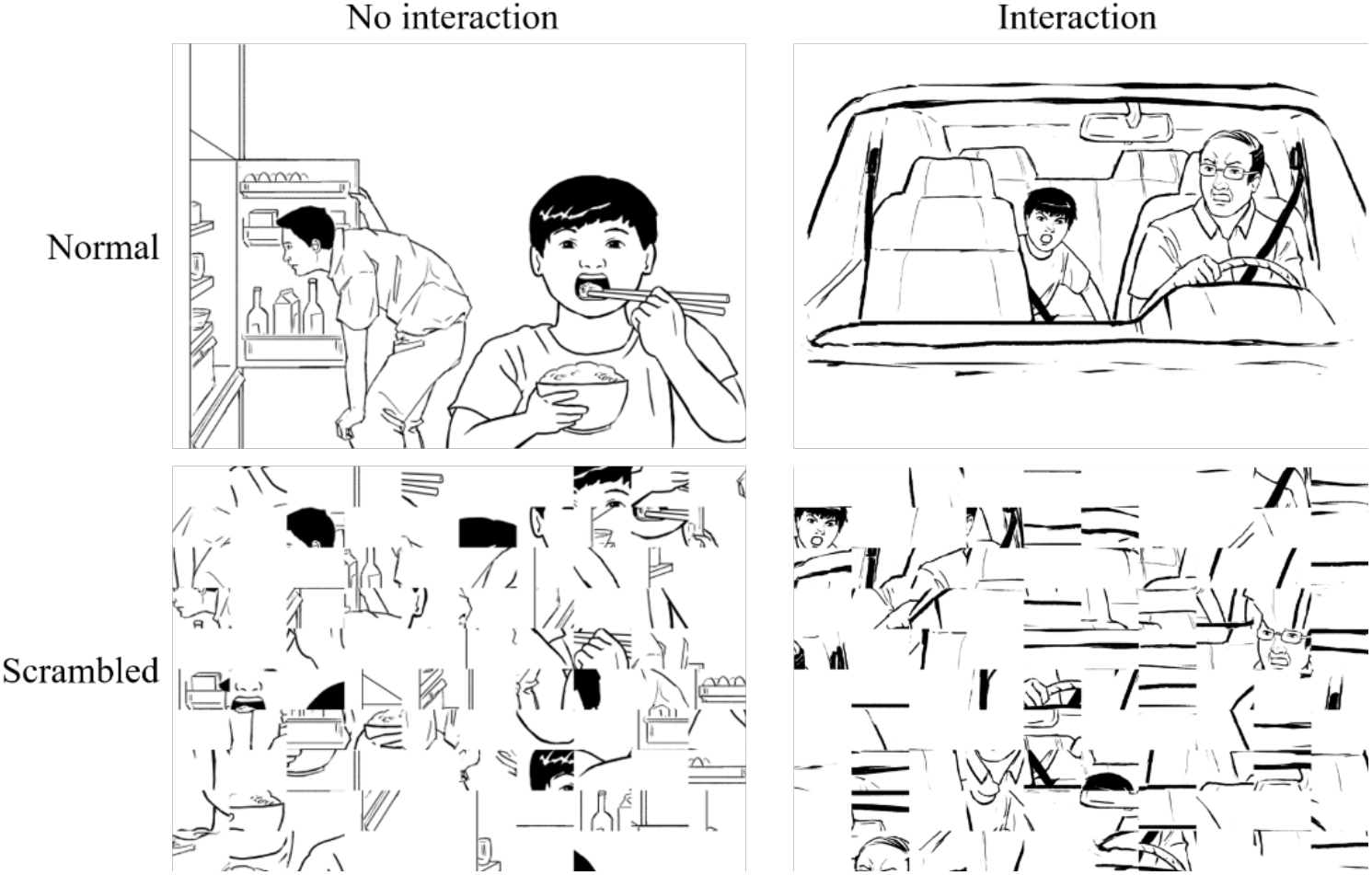
One example image of the four stimulus types (Oomen, 2022).

**Figure 2.**
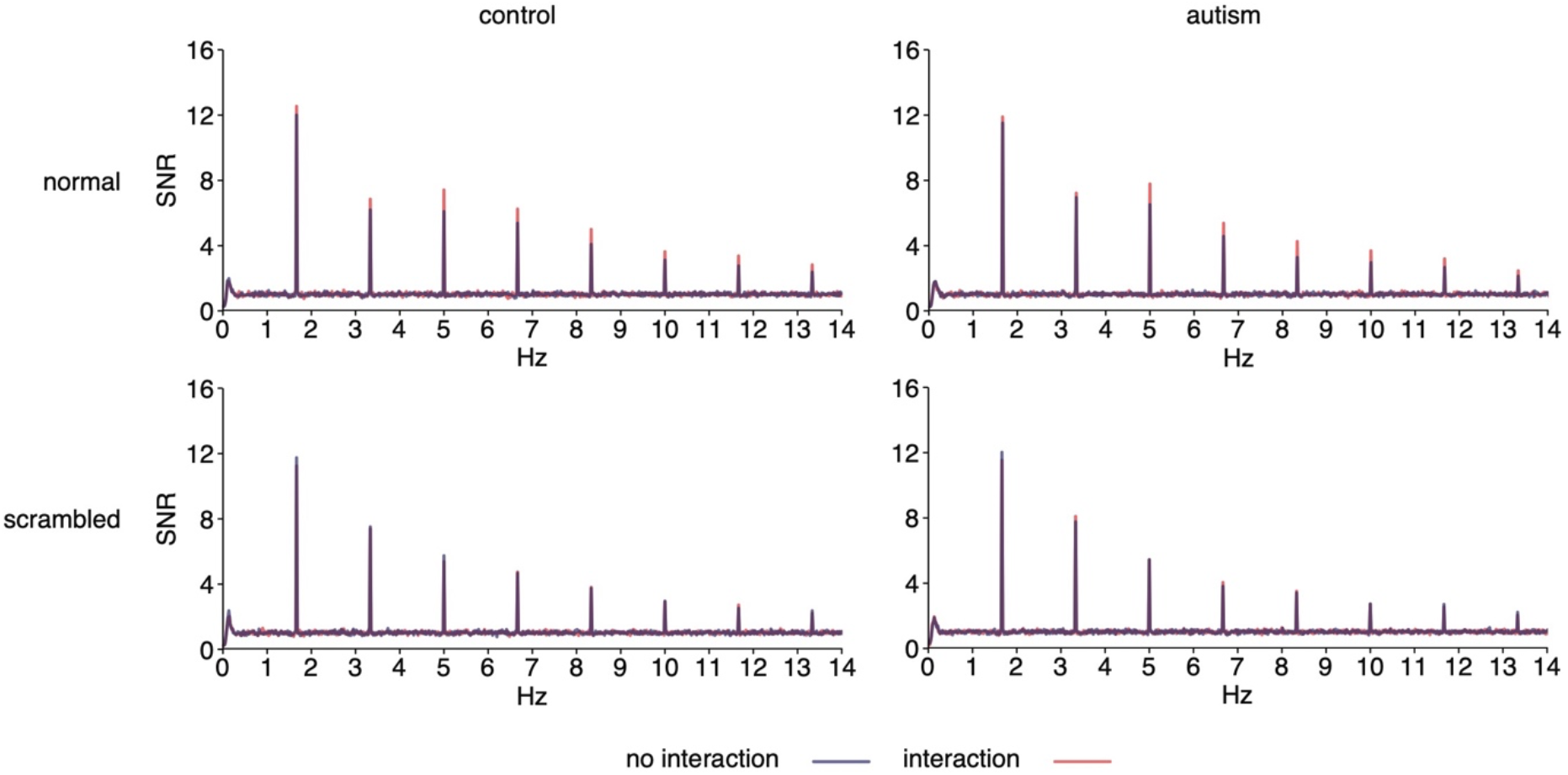
Signal-to-noise (SNR) over electrodes of interest per condition per group.

**Figure 3.**
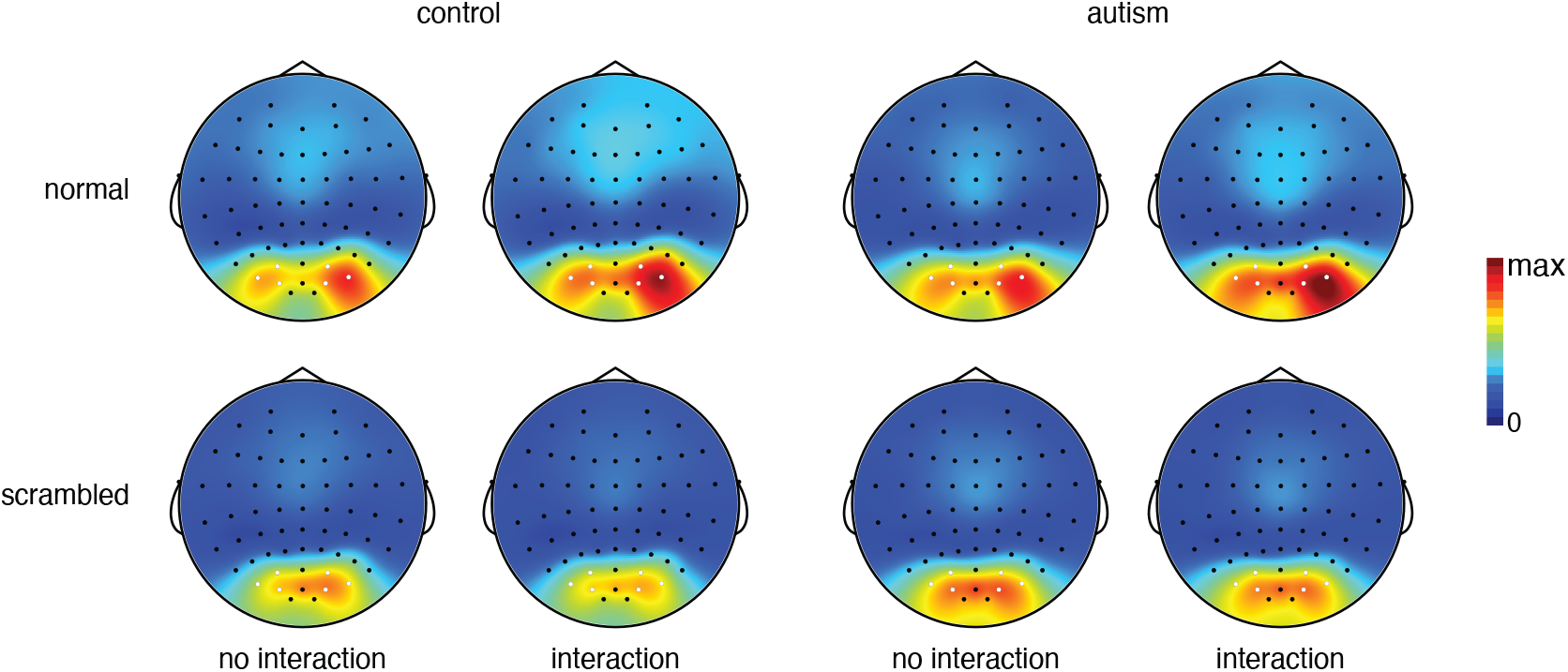
Topographies per group, per condition. Topographies are scaled from 0 to the maximum amplitude of the four conditions and two groups (i.e. 5.19 µV). Included electrodes are indicated in white.

In addition to the confirmatory analyses reported above, we also ran explorative (but preregistered) correlation analyses investigating whether social interaction processing correlated with autism symptomatology (SRS-A and AQ) and alexithymia (TAS-20). These correlations were not significant with Bayesian analyses revealing moderate evidence against a correlation between social interaction processing and autism symptomatology (SRS-A: *r*_s_ = - 0.03, *p* = .842, BF_10_ = 0.29; AQ: *r*_s_ = 0.04, *p* = .823, BF_10_ = 0.29) or alexithymia (TAS-20: *r*_s_ = -0.09, *p* = .474, BF_10_ = 0.31).

## 4. Discussion

To explain the social difficulties in autism, a large body of neuroscience research has studied social stimuli processing (e.g. a single agent: Nijhof et al., 2018; or mere body parts: Kang et al., 2018). In contrast, there are, to the best of our knowledge, no neuroscience studies that have investigated social interaction processing. Existing behavioural studies point to altered social interaction processing in autism, however it is not clear whether this can be attributed to the fundamental process of social interaction recognition, which we tested in the current study. If the fundamental process of recognizing social interaction is anomalous in autism, this could have a cascading effect on how people with autism extract information from, and in turn adapt their behaviour to, third-party social interactions. We therefore applied an EEG frequency tagging task that was previously validated by Oomen et al. (2022) to study social interaction recognition. More precisely, we investigated the process of inferring social interaction from context by measuring neural responses to social scenes depicting either social interaction or not in adults with and without autism. As hypothesized, we replicated the findings by Oomen et al. (2022), that is, we found stronger responses over predominantly right-lateralized occipitoparietal electrodes to social scenes depicting two interacting agents than to social scenes depicting two non-interacting agents. However, contrary to our expectations, we found this effect in both groups with no difference between them.

By replicating the findings of Oomen et al. (2022), the current study increases confidence in the reliability of the EEG frequency task used to measure social interaction recognition objectively. Not only is this important for the current study, but also, more broadly, for possible future studies into social interaction recognition that aim to answer more fundamental questions (e.g. how valence modulates social interaction recognition) or questions related to other clinical conditions that are associated with social difficulties (e.g. Williams syndrome, Schizophrenia, Social anxiety, or Personality Disorders; Kennedy & Adolphs, 2012). Besides replicating the previously found social interaction recognition effect (Oomen et al., 2022) in our control group, we also found the same effect in the autism group, with no difference between the groups. This lack of group difference was supported by a Bayesian test, and by a correlational analysis that showed no relationship between social interaction processing and autism symptomatology. Thus, our results show that social interaction recognition is not anomalous in autism and can therefore not explain the social difficulties people with autism experience.

Our findings may seem at odds with previous behavioural studies that suggest aberrant processing of social interactions in autism (e.g. Liu et al., 2018; van Boxtel et al., 2017; von der Lühe et al., 2016). However, as aforementioned, these behavioural results can be explained in two ways: 1) Individuals with autism have difficulties to recognize social interaction or 2) individuals with autism are perfectly able to recognize social interactions, but do not extract the same information from those interactions and/or use this information similarly. The current study measured social interaction processing objectively, that is, without an explicit task, and found no differences between groups. As such, it dismisses the first explanation and helps interpret the findings of previous behavioural research on social interaction processing and autism. For example, Liu et al. (2018) showed that motion sequences that depict agents in social interaction tend to be perceived as shorter in duration by neurotypicals, but that this effect was negatively correlated with autism symptomatology.Following the above reasoning, the results of the current study suggest that this negative correlation may not be due to atypical social interaction recognition per se, but due to something else that affects a more cognitive process of perception of time (e.g. enjoyability: Agarwal & Karahanna (2000); arousal: Gil & Droit-Volet, 2011).

Similarly, von der Lühe et al. (2016) found that adults with autism show diminished interpersonal predictive coding. Here again, following our above reasoning, this findings by von der Lühe et al. (2016) might indicate that, although individuals with autism spontaneously recognize social interactions, they take less advantage of the acquired social information to anticipate other’s actions (i.e. the interactive actions of one agent to predict the response of the interaction partner). Lastly, in a neurotypical sample, van Boxtel et al. (2017) found that adults who score relatively high on autism symptomatology showed a reduced ability to differentiate between interactive and non-interactive actions. Furthermore, they found a negative correlation between autism symptomatology and interactivity ratings for the interactive actions. However, these results are acquired by an explicit task in which participants had to judge the degree of interaction. As a result, it might be that autism is not associated with a reduced ability to recognize social interactions, but that individuals who score relatively high on autism happen to judge the degree of social interaction differently. That is, one can judge a social interaction as more or less engaging, while still detecting that there is in fact social interaction. Put more generally, our results suggest that it is not social interaction recognition per se that is anomalous in autism, but rather how individuals with autism interpret and act upon the information they extract from such interactions. An important avenue for future research will be to further explore this hypothesis and specifically to investigate the degree to which anomalies in extracting information from and responding to social interactions can explain social difficulties in autism.

Besides an enhanced neural response to interactive (vs. non-interactive) normal stimuli for both the autism and the control group, we also found a difference between interactive and non-interactive scrambled stimuli, but in the opposite direction. More specifically, we found an enhanced response for the scrambled stimuli without social interaction compared to the scrambled stimuli with social interaction. Although this reversed pattern can be observed in the original study as well (Oomen et al., 2022), the difference was not significant there, possibly due to the smaller sample (*N* = 28 versus 61 in the current study). Differences in the neural response elicited by scrambled stimuli suggest the presence of low-level differences (e.g. luminance) influencing the brain response. Crucially, however, these low-level influences acted in the opposite direction as the influence of social interaction. Therefore, they cannot explain the observed enhancement of neural responses for interaction (vs. non-interaction) normal stimuli. If anything, the opposite effect for scrambled stimuli indicates that the strength of the effect in the normal stimuli may have been slightly underestimated and further highlights the importance of comparing the experimental effects to a baseline condition capturing low-level differences.

This study has two limitations that can be addressed by future research. First, our results are limited to an autism population that matches our sample characteristics (e.g. adults with an IQ above 85). We therefore cannot generalize our findings to other individuals on the autism spectrum, such as children or those with a lower IQ. Future studies are warranted to test the degree to which our findings generalize to such populations, as well as the developmental trajectory of social interaction recognition. In fact, the task used here lends itself well for such research, as it does not require verbal task instructions and can measure brain responses with high precision, both of which are important advantages for research in younger populations (infants, and young kids), who have limited verbal capabilities and often have difficulties to sit still (Azhari et al., 2020; Maguire et al., 2014; Raschle et al., 2012).

A second limitation is that participants were aware of the stimulus categories. In line with the original study (Oomen et al., 2022), we were open about the included stimulus categories during the introduction of the task. Because block designs allow participants to become aware of the categories regardless, providing the participants with this information beforehand reduces variability regarding the timepoint at which participants recognize the stimulus categories. It is important to note however, that awareness of the stimuli categories is by no means unique to our study design. Previous autism (ERP or frequency tagging) studies were often either open about the stimulus categories (own vs close-other name or face, e.g. Cygan et al., 2014; Nijhof et al., 2018; Nowicka et al., 2016) or the stimulus categories were easily identifiable (face vs. object: Sysoeva et al. (2018); face vs. house: Vettori et al., 2020). Furthermore, although we were open about the stimulus categories to the participants, the tasks they had to perform were not connected to the stimulus categories, and participants were not aware of the research question. Nevertheless, it would be interesting for future research to investigate the role of prior awareness of the stimulus categories on brain responses.

To conclude, we found an enhanced response to social scenes with social interaction compared to social scenes without social interaction over predominantly right-lateralized occipitoparietal electrodes, replicating previous findings in a neurotypical population (Oomen et al., 2022). However, we found this effect in the control group as well as the autism group with no difference between them. This suggests that social interaction recognition is not anomalous in adults with autism and cannot explain their social difficulties experienced in daily life.

## Acknowledgements

The authors would like to thank all participants for their contribution as well as Amber Konings, Delphine Libeer, and Joyce Scheirlinckx for their assistance with data collection and recruitment.

## Supplementary material

**Figure 1.**
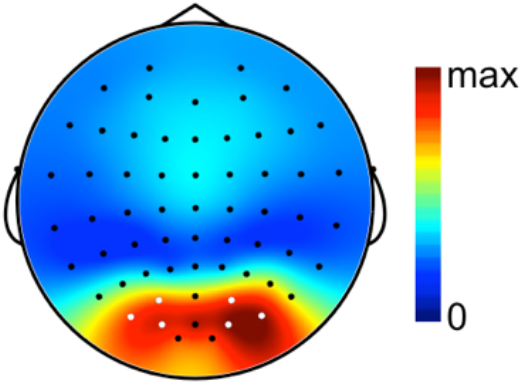
Topography across groups and conditions, scaled from 0 to maximum amplitude (i.e. 4.14 µV)

## Footnotes

1 We acknowledge and respect different preferences for language used to refer to a person with a diagnosis of ASD. Here, we use ‘person with autism’ to respect the preference of the autism sample described in this study.

2 This included ASD (DSM-5) as well as diagnoses based on earlier DSM versions, such as Autistic Disorder, Asperger’s disorder, and Pervasive Developmental Disorder—Not Otherwise Specified.

## Literature

Agarwal, R., & Karahanna, E. (2000). Time flies when you’re having fun. MIS Quarterly, 24(4), 665–694. https://doi.org/10.2307/3250951

American Psychiatric Association. (2013). DSM-5 Diagnostic Classification. In Diagnostic and Statistical Manual of Mental Disorders. https://doi.org/10.1176/appi.books.9780890425596.x00diagnosticclassification

Azhari, A., Truzzi, A., Neoh, M. J. Y., Balagtas, J. P. M., Tan, H. A. H., Goh, P. L. P., Ang, X. H. A., Setoh, P., Rigo, P., Bornstein, M. H., & Esposito, G. (2020). A decade of infant neuroimaging research: What have we learned and where are we going? Infant Behavior and Development, 58, 101389. https://doi.org/10.1016/J.INFBEH.2019.101389

Bagby, R. M., Parker, J. D. A., & Taylor, G. J. (1994). The twenty-item Toronto Alexithymia scale—I. Item selection and cross-validation of the factor structure. Journal of Psychosomatic Research, 38(1), 23–32. https://www.academia.edu/13022526/The_twenty_item_Toronto_Alexithymia_scale_I_Item_selection_and_cross_validation_of_the_factor_structure

Baron-Cohen, S., Wheelwright, S., Skinner, R., Martin, J., & Clubley, E. (2001). The autism-spectrum quotient (AQ): evidence from Asperger syndrome/high-functioning autism, males and females, scientists and mathematicians. Journal of Autism and Developmental Disorders, 31(1), 5–17. https://doi.org/10.1023/A:1005653411471

Christ, O., Schmid, K., Lolliot, S., Swart, H., Stolle, D., Tausch, N., Ramiah, A. al, Wagner, U., Vertovec, S., & Hewstone, M. (2014). Contextual effect of positive intergroup contact on outgroup prejudice. Proceedings of the National Academy of Sciences of the United States of America, 111(11), 3996–4000. https://doi.org/10.1073/pnas.1320901111

Constantino, J. N. (2002). The social responsiveness scale. Western psychological services.

Costanzo, M., & Archer, D. (1989). Interperting the expressive behavior of others: The Interpersonal Perception Task. Journal of Nonverbal Behavior, 13(4), 225–245. https://doi.org/10.1007/BF00990295

Cowell, J. M., & Decety, J. (2015). Precursors to morality in development as a complex interplay between neural, socioenvironmental, and behavioral facets. Proceedings of the National Academy of Sciences of the United States of America, 112(41), 12657–12662. https://doi.org/10.1073/pnas.1508832112

Cygan, H. B., Tacikowski, P., Ostaszewski, P., Chojnicka, I., & Nowicka, A. (2014). Neural correlates of own name and own face detection in autism spectrum disorder. PLoS ONE, 9(1). https://doi.org/10.1371/journal.pone.0086020

Dziobek, I., Fleck, S., Kalbe, E., Rogers, K., Hassenstab, J., Brand, M., Kessler, J., Woike, J. K., Wolf, O. T., & Convit, A. (2006). Introducing MASC: a movie for the assessment of social cognition. Journal of Autism and Developmental Disorders, 36(5), 623–636. https://doi.org/10.1007/S10803-006-0107-0

Efran, M. G., & Cheyne, J. A. (1973). Shared space: The co-operative control of spatial areas by two interacting individuals. Canadian Journal of Behavioural Science/Revue Canadienne Des Sciences Du Comportement, 5(3), 201–210. https://doi.org/10.1037/h0082345

Gil, S., & Droit-Volet, S. (2011). Time perception in response to ashamed faces in children and adults. Scandinavian Journal of Psychology, 52(2), 138–145. https://doi.org/10.1111/J.1467-9450.2010.00858.X

Golan, O., Baron-Cohen, S., & Golan, Y. (2008). The “Reading the Mind in Films” Task [child version]: complex emotion and mental state recognition in children with and without autism spectrum conditions. Journal of Autism and Developmental Disorders, 38(8), 1534–1541. https://doi.org/10.1007/S10803-007-0533-7

Goris, J., Braem, S., Nijhof, A. D., Rigoni, D., Deschrijver, E., van de Cruys, S., Wiersema, J. R., & Brass, M. (2018). Sensory Prediction Errors Are Less Modulated by Global Context in Autism Spectrum Disorder. Biological Psychiatry: Cognitive Neuroscience and Neuroimaging, 3(8), 667–674. https://doi.org/10.1016/J.BPSC.2018.02.003

Goris, J., Braem, S., van Herck, S., Simoens, J., Deschrijver, E., Wiersema, J. R., Paton, B., Brass, M., & Todd, J. (2022). Reduced Primacy Bias in Autism during Early Sensory Processing. Journal of Neuroscience, 42(19), 3989–3999. https://doi.org/10.1523/JNEUROSCI.3088-20.2022

Heavey, L., Phillips, W., Baron-Cohen, S., & Rutter, M. (2000). The Awkward Moments Test: a naturalistic measure of social understanding in autism. Journal of Autism and Developmental Disorders, 30(3), 225–236. https://doi.org/10.1023/A:1005544518785

Hoekstra, R. A., Bartels, M., Cath, D. C., & Boomsma, D. I. (2008). Factor structure, reliability and criterion validity of the Autism-Spectrum Quotient (AQ): a study in Dutch population and patient groups. Journal of Autism and Developmental Disorders, 38(8), 1555–1566. https://doi.org/10.1007/S10803-008-0538-X

Isik, L., Mynick, A., Pantazis, D., & Kanwisher, N. (2020). The speed of human social interaction perception. NeuroImage, 215, 116844. https://doi.org/10.1016/j.neuroimage.2020.116844

Kang, E., Keifer, C. M., Levy, E. J., Foss-Feig, J. H., McPartland, J. C., & Lerner, M. D. (2018). Atypicality of the N170 Event-Related Potential in Autism Spectrum Disorder: A Meta-analysis. Biological Psychiatry: Cognitive Neuroscience and Neuroimaging, 3(8), 657–666. https://doi.org/10.1016/j.bpsc.2017.11.003

Kennedy, D. P., & Adolphs, R. (2012). The social brain in psychiatric and neurological disorders. Trends in Cognitive Sciences, 16(11), 559. https://doi.org/10.1016/J.TICS.2012.09.006

Kinnaird, E., Stewart, C., & Tchanturia, K. (2019). Investigating alexithymia in autism: A systematic review and meta-analysis. European Psychiatry, 55, 80. https://doi.org/10.1016/J.EURPSY.2018.09.004

Knowles, E. S. (2015). Spatial behavior of individuals and groups. In Psychology of group influence (2nd ed., pp. 53–86). Psychology Press.

Liu, R., Yuan, X., Chen, K., Jiang, Y., & Zhou, W. (2018). Perception of social interaction compresses subjective duration in an oxytocin-dependent manner. ELife, 7. https://doi.org/10.7554/ELIFE.32100

Luck, S. J., & Gaspelin, N. (2017). How to Get Statistically Significant Effects in Any ERP Experiment (and Why You Shouldn’t). Psychophysiology, 54(1), 146. https://doi.org/10.1111/PSYP.12639

Maguire, M. J., Magnon, G., & Fitzhugh, A. E. (2014). Improving data retention in EEG research with children using child-centered eye tracking. Journal of Neuroscience Methods, 238, 78. https://doi.org/10.1016/J.JNEUMETH.2014.09.014

Mast, M. S., & Hall, J. A. (2004). Who is the boss and who is not? Accuracy of judging status. Journal of Nonverbal Behavior, 28(3), 145–165. https://doi.org/10.1023/B:JONB.0000039647.94190.21

Morey, R. D. (2008). Confidence Intervals from Normalized Data: A correction to Cousineau (2005). Tutorial in Quantitative Methods for Spsychology, 4(2), 61–64.

Nijhof, A. D., Bardi, L., Brass, M., & Wiersema, J. R. (2018). Brain activity for spontaneous and explicit mentalizing in adults with autism spectrum disorder: An fMRI study. NeuroImage: Clinical, 18, 475–484. https://doi.org/10.1016/J.NICL.2018.02.016

Nijhof, A. D., Dhar, M., Goris, J., Brass, M., & Wiersema, J. R. (2018). Atypical neural responding to hearing one’s own name in adults with ASD. Journal of Abnormal Psychology, 127(1), 129–138. https://doi.org/10.1037/abn0000329

Noens, I., de la Marche, W., & Scholte, E. (2012). Screeningslijst voor autismespectrumstoornissen. Hogrefe Ultgevers B.V.

Norcia, A. M., Gregory Appelbaum, L., Ales, J. M., Cottereau, B. R., & Rossion, B. (2015a). The steady-state visual evoked potential in vision research: A review. Journal of Vision, 15(6), 1–46. https://doi.org/10.1167/15.6.4

Norcia, A. M., Gregory Appelbaum, L., Ales, J. M., Cottereau, B. R., & Rossion, B. (2015b). The steady-state visual evoked potential in vision research: A review. Journal of Vision, 15(6), 1–46. https://doi.org/10.1167/15.6.4

Nowicka, A., Cygan, H. B., Tacikowski, P., Ostaszewski, P., & Kuś, R. (2016). Name recognition in autism: EEG evidence of altered patterns of brain activity and connectivity. Molecular Autism, 7(1), 1–14. https://doi.org/10.1186/s13229-016-0102-z

Oomen, D., Cracco, E., Brass, M., & Wiersema, J. R. (2022). EEG frequency tagging evidence of social interaction recognition. Social Cognitive and Affective Neuroscience, 00, 1–10. https://doi.org/10.1093/SCAN/NSAC032

Peirce, J., Gray, J. R., Simpson, S., MacAskill, M., Höchenberger, R., Sogo, H., Kastman, E., & Lindeløv, J. K. (2019). PsychoPy2: Experiments in behavior made easy. Behavior Research Methods, 51(1), 195–203. https://doi.org/10.3758/s13428-018-01193-y

Quadflieg, S., & Koldewyn, K. (2017). The neuroscience of people watching: how the human brain makes sense of other people’s encounters. Annals of the New York Academy of Sciences, 1396(1), 166–182. https://doi.org/10.1111/NYAS.13331

Quadflieg, S., & Penton-Voak, I. S. (2017). The Emerging Science of People-Watching: Forming Impressions From Third-Party Encounters. Current Directions in Psychological Science, 26(4), 383–389. https://doi.org/10.1177/0963721417694353

Raschle, N., Zuk, J., Ortiz-Mantilla, S., Sliva, D. D., Franceschi, A., Grant, P. E., Benasich, A., & Gaab, N. (2012). Pediatric neuroimaging in early childhood and infancy: challenges and practical guidelines. Annals of the New York Academy of Sciences, 1252(1), 43. https://doi.org/10.1111/J.1749-6632.2012.06457.X

Raymaekers, R., Wiersema, J. R., & Roeyers, H. (2009). EEG study of the mirror neuron system in children with high functioning autism. Brain Research, 1304, 113–121. https://doi.org/10.1016/J.BRAINRES.2009.09.068

Retter, T. L., & Rossion, B. (2016). Uncovering the neural magnitude and spatio-temporal dynamics of natural image categorization in a fast visual stream. Neuropsychologia, 91, 9–28. https://doi.org/10.1016/j.neuropsychologia.2016.07.028

Retter, T. L., Rossion, B., & Schiltz, C. (2021). Harmonic Amplitude Summation for Frequency-tagging Analysis. Journal of Cognitive Neuroscience, 33(11), 2372–2393. https://doi.org/10.1162/JOCN_A_01763

Roeyers, H., Buysse, A., Ponnet, K., & Pichal, B. (2001). Advancing Advanced Mind-reading Tests: Empathic Accuracy in Adults with a Pervasive Developmental Disorder. Journal of Child Psychology and Psychiatry, 42(2), 271–278. https://doi.org/10.1111/1469-7610.00718

Rouder, J. N., Morey, R. D., Speckman, P. L., & Province, J. M. (2012). Default Bayes factors for ANOVA designs. Journal of Mathematical Psychology, 56(5), 356–374. https://doi.org/10.1016/J.JMP.2012.08.001

Sinke, C. B. A., Sorger, B., Goebel, R., & de Gelder, B. (2010). Tease or threat? Judging social interactions from bodily expressions. NeuroImage, 49(2), 1717–1727. https://doi.org/10.1016/j.neuroimage.2009.09.065

Sysoeva, O. v., Constantino, J. N., & Anokhin, A. P. (2018). Event-related potential (ERP) correlates of face processing in verbal children with autism spectrum disorders (ASD) and their first-degree relatives: a family study. Molecular Autism, 9(1). https://doi.org/10.1186/S13229-018-0220-X

Trijsburg, R. W., Passchier, J., Duivenvoorden, H., & Bagby, R. M. (1997). The Toronto Alexithymia Scale (Dutch version). Department of Medical Psychology and Psychotherapy, Erasmus University.

van Boxtel, J. J. A., Peng, Y., Su, J., & Lu, H. (2017). Individual differences in high-level biological motion tasks correlate with autistic traits. Vision Research, 141, 136–144. https://doi.org/10.1016/J.VISRES.2016.11.005

Vettori, S., Dzhelyova, M., van der Donck, S., Jacques, C., Steyaert, J., Rossion, B., & Boets, (2019). Reduced neural sensitivity to rapid individual face discrimination in autism spectrum disorder. NeuroImage: Clinical, 21, 101613. https://doi.org/10.1016/J.NICL.2018.101613

Vettori, S., Dzhelyova, M., van der Donck, S., Jacques, C., van Wesemael, T., Steyaert, J., Rossion, B., & Boets, B. (2020). Combined frequency-tagging EEG and eye tracking reveal reduced social bias in boys with autism spectrum disorder. Cortex; a Journal Devoted to the Study of the Nervous System and Behavior, 125, 135–148. https://doi.org/10.1016/J.CORTEX.2019.12.013

von der Lühe, T., Manera, V., Barisic, I., Becchio, C., Vogeley, K., & Schilbach, L. (2016). Interpersonal predictive coding, not action perception, is impaired in autism. Philosophical Transactions of the Royal Society of London. Series B, Biological Sciences, 371(1693). https://doi.org/10.1098/RSTB.2015.0373

Wechsler, D. (2008). Wechsler Adult Intelligence Scale–Fourth Edition (WAIS-IV). NCS Pearson .

